# NMDA spikes mediate amplification of odor pathway information in the piriform cortex

**DOI:** 10.1101/346791

**Authors:** Amit Kumar, Oded Schiff, Edi Barkai, Bartlett W. Mel, Alon Poleg-Polsky, Jackie Schiller

**Affiliations:** Department of Physiology, The Rappaport Faculty of Medicine and Research Institute, Technion-Israel Institute of Technology, Haifa, Israel.; Department of Physiology and Biophysics, University of Colorado School of Medicine, Aurora, CO, USA; University of Haifa, Department of Neurobiology, Haifa, Israel; Biomedical Engineering Department, University of Southern California, Los Angeles, CA, USA

## Abstract

The piriform cortex (PCx) receives direct input from the olfactory bulb (OB) and is the brain’s main station for odor recognition and memory. The transformation of the odor code from OB to PCx is profound: mitral and tufted cells in olfactory glomeruli respond to individual odorant molecules, whereas pyramidal neurons (PNs) in the PCx responds to multiple, apparently random combinations of activated glomeruli. How these “discontinuous” receptive fields are formed from OB inputs remains unknown. Counter to the prevailing view that olfactory PNs sum their inputs passively, we show for the first time that NMDA spikes within individual dendrites can both amplify OB inputs and impose combination selectivity upon them, while their ability to compartmentalize voltage signals allows different dendrites to represent different odorant combinations. Thus, the 2-layer integrative behavior of olfactory PN dendrites provides a parsimonious account for the nonlinear remapping of the odor code from bulb to cortex.

## Introduction

The piriform cortex (PCx) is the main cortical station in olfactory processing. It receives direct odor information from the olfactory bulb, as well as contextual information from higher brain regions, and is thought to be the brain’s primary site for odor discrimination and recognition ^1^.

Olfaction starts at the nasal epithelium where a single odor activates multiple odorant receptors (ORs). At the olfactory bulb, information from like ORs converge to ∼1000 mirror symmetric pairs of glomeruli ^2,3^. Thus, on either side of the brain an odor is represented as a distributed pattern of activation over ∼1000 glomeruli, which together form a molecular map of the odor. Mitral and tufted (M/T) cells, the glomerular outputs, carry the olfactory signal next to the PCx via the lateral olfactory tract (LOT). LOT axons traverse layer 1 of the PCx and form synaptic contacts with the thin distal dendrites of pyramidal neurons and layer 1 interneurons ^1,4^.

Unlike other sensory cortices which are topographically organized, the connectivity scheme between the OB and PCx lacks any apparent spatial structure: individual LOT axons from the bulb terminate in broad overlapping swaths of the PCx ^5-8^ so that each glomerulus excites a widely dispersed and apparently randomly distributed population of PCx neurons. In turn, each pyramidal neuron in PCx receives synaptic input from roughly ∼10% of the 1000 glomeruli, also apparently randomly sampled ^8-14^. In keeping with the notion of random convergence and divergence of OB axons in PCx, electrophysiological studies show that each odorant activates an apparently random population of from 3% to 15% of the neurons in layer 2 of the PCx at low concentration ^13^.

Pyramidal neurons in the piriform cortex are the main integration units within which the discrete molecular information channels of the OB are combined to form “odor objects”, but the biophysical and circuit-level mechanisms that remap LOT inputs into the olfactory code in PCx remain poorly understood. Three features of the mapping of LOT excitation into pyramidal neuron activity in PCx are noteworthy, and in the context of the existing literature, lead to a conundrum:

1. Pyramidal neurons are reliably driven by LOT inputs, even though LOT synaptic contacts onto PNs are formed on distal tuft dendrites, are few in number (∼200 total contacts, ^8^), and are sparsely activated (just 1% of glomeruli activated in the OB reliably drives many PCx pyramidal neurons ^9^. This suggests pyramidal neurons in PCx have some means of amplifying weak distal inputs.
2. Pyramidal neurons in PCx are combination selective, that is, they respond supralinearly to specific combinations of glomerular inputs but not others ^9,15^. These combinations are of relatively high order, so that a pyramidal neuron that responds strongly to odorant may respond weakly to a chemically similar odorant that activates a heavily overlapping pattern of glomeruli ^13^.
3. Pyramidal neurons in PCx have “discontinuous” receptive fields, that is, they respond to multiple chemically diverse odorants ^13,16-18^, while failing to respond to re-combinations of those odorants’ component parts ^9,15^. For example, a pyramidal neuron that is unresponsive to individual odor components A, B, or C may respond strongly to combinations AB and AC but fail to respond to combination BC.

What biophysical mechanism(s) could account for a pyramidal neuron’s ability to (1) amplify distal LOT inputs; (2) enforce combination selectivity on those inputs; and (3) maintain multiple “discontinuous” recognition subunits? A possible mechanism could be compartmentalized NMDA spikes in pyramidal neuron dendrites, which could provide both, the thresholding nonlinearity that enforces combination selectivity, and the amplification that allows distal inputs to drive somatic action potentials ^19-25^.

Weighing against this hypothesis, however, the one study that has analyzed dendritic responses of olfactory PNs using current injections and focal electrical stimulation reported that pyramidal neurons in the PCx lack sufficient NMDA (or other) regenerative currents that could provide either the combination selectivity or amplification of LOT inputs ^26^. Rather, they reported that pyramidal neurons in PCx, unlike their counterparts in other cortical areas, act as intrinsically linear summing units. This leads to a conundrum: the only alternative source of nonlinearity that would seem capable of producing combination selectivity – recurrent network effects mediated by intracortical inputs (IC) to pyramidal neurons – has also apparently been ruled out: Davison and Ehlers (2009) tested whether a pyramidal neuron’s ability to respond selectively to LOT input combinations depends on IC inputs, which outnumber a pyramidal neuron’s LOT inputs 10 to 1, but they found no reduction in a pyramidal neuron’s combination selectively when its IC inputs were blocked with baclofen.

Given the importance of understanding the cellular mechanisms underlying odor representation in PCx. we revisited the question as to whether pyramidal neuron dendrites in PCx can generate local spikes ^21,24,27-31^. We found that robust NMDA spikes can indeed be generated in dendrites of PCx pyramidal neurons both in layer 1a which receives direct LOT input, as well as in deeper layers, and using a model we show that these local spikes can serve to effectively amplify clustered versus distributed LOT inputs forming the basis for a discontinuous receptive field. We also show that supralinear summation of LOT inputs is largely confined to a single dendrite whereas nonlinear interactions of LOT inputs between dendrites, are small. These findings support the idea that a pyramidal neuron in PCx can represent multiple distinct glomerular combinations within its apical dendritic arbor, which fulfills the basic requirements for a discontinuous receptive field ^13^. Finally we show that interactions between LOT and IC inputs are also nonlinear, a fact that will likely be important for understanding the recurrent pattern completion functions of the PCx.

## Results

### Glutamate uncaging evoked NMDA spikes in apical dendrites of PCx pyramidal neurons

To directly address the fundamental question of whether dendrites of pyramidal neurons in PCx can generate dendritic spikes we used focal glutamate uncaging (MNI-glutamate) to activate specific dendritic locations. Neurons were loaded with the calcium sensitive dye OBG-6F (200 µM) and CF633 (200 µM) to visualize the dendritic tree and perform calcium imaging (Figure 1a) Gradually increasing the laser intensity to uncage glutamate, evoked EPSP-like potentials, which increased progressively up to a threshold laser activation, beyond which a local spike was initiated (Figure 1a-b; n=10 cells). We then perfused the slices with specific blockers for voltage-gated sodium, calcium and NMDAR channels ^28,32^. Addition of the voltage-gated sodium channel blocker TTX (1 µm) did not significantly change the peak of the slow spikes (90.8 ± 60 % of control; p=0.16; n=5) In some cases, we recorded a fast spikelet that preceded the prolonged voltage plateau, reminiscent of the sodium spikelets observed in CA1 pyramidal dendrites and tuft dendrites of neocortical layer 5 pyramidal neurons ^21,33^; we verified that the fast spikelet was blocked by TTX (example of the fast spikelet is shown in Figure 2f and Figure 4b). An add-on cocktail of voltage-gated calcium channel blockers (w-agatoxin 0.5 µM, conotoxin-GVIA 5 µM; SNX 482 200 nM, nifedipine 10µM) had a small effect on the peak of the dendritic spike (85.4 ± 4.2 % of control; p=0.08; n=5). Finally, APV (2-amino 5-phosphonovalerate; 50 µM) a specific NMDAR channel blocker, completely abolished spike initiation (Figure 1b-d). These results indicate that like fine dendrites of hippocampal and neocortical pyramidal neurons ^31,34^, NMDA spikes can be initiated in dendrites of pyramidal neurons in PCx. Voltage-gated sodium and calcium channels only minimally contributed to the slow component of the spike, while the majority of the current was carried by NMDAR channels ^28,32^.

**Figure 1.**
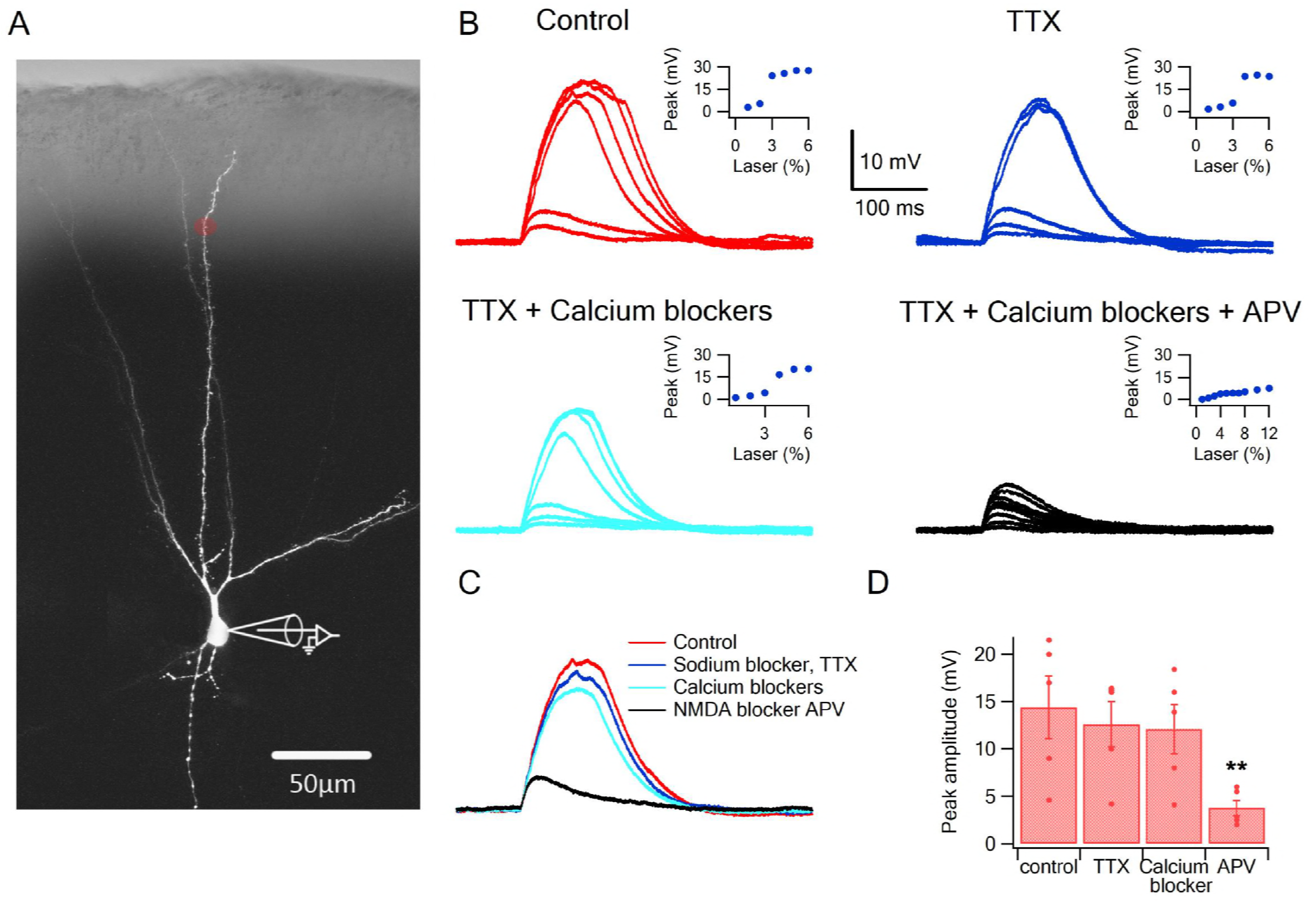
Glutamate uncaging evoked NMDA spikes in dendrites of PCx pyramidal neurons. a. Fluorescence image reconstruction of a pyramidal neuron filled with CF633 (200 µM) via the patch recording electrode. Uncaging location is indicated by the red dot. b. Voltage responses and dendritic spikes were evoked by uncaging of MNI-glutamate at increasing laser intensities in the Control condition (red); in the presence of the voltage gated sodium channel blocker TTX (1 µm; blue); with an additional cocktail of voltage-gated calcium channel blockers (w-agatoxin 0.5 µM, conotoxin-GVIA 5 µM; SNX 482 200 nM, nifedipine 10µM; cyan), and finally adding APV (50 µm; black). Insets, peak voltage response at increasing laser intensity for the different conditions. c. Overlay of the spike in the Control condition (red), in TTX (blue), in TTX+ calcium channel blockers (cyan) and with APV (black). All traces were collected using the same laser stimulation intensity. The spike was completely blocked with APV, and could not be reinitiated at higher laser intensities. d. Summary plot of the peak response amplitude in control, TTX, Ca^2+^ blockers, and APV (n=5). ** p<0.01 for comparison of APV with other blockers.

### Initiation of NMDA spikes by activation of LOT inputs using focal synaptic stimulation

Having established with focal glutamate uncaging that pyramidal neurons in PCx are capable of producing NMDA spikes in their distal dendrites, we asked whether synaptic activation of LOT inputs can also trigger dendritic spikes. We directly activated LOT inputs using synaptic stimulation electrodes visually positioned within the LOT pathway in layer 1 ^15,35^. (Figure 2a). The dendritic stimulus location was verified using calcium imaging which showed a low amplitude localized calcium transient in response to a small subthreshold EPSP (Figure 2b). Gradually increasing the stimulus intensity led to a linearly increasing EPSP up to a threshold stimulation intensity, after which a dendritic spike was initiated (Figure 2c-d).

**Figure 2.**
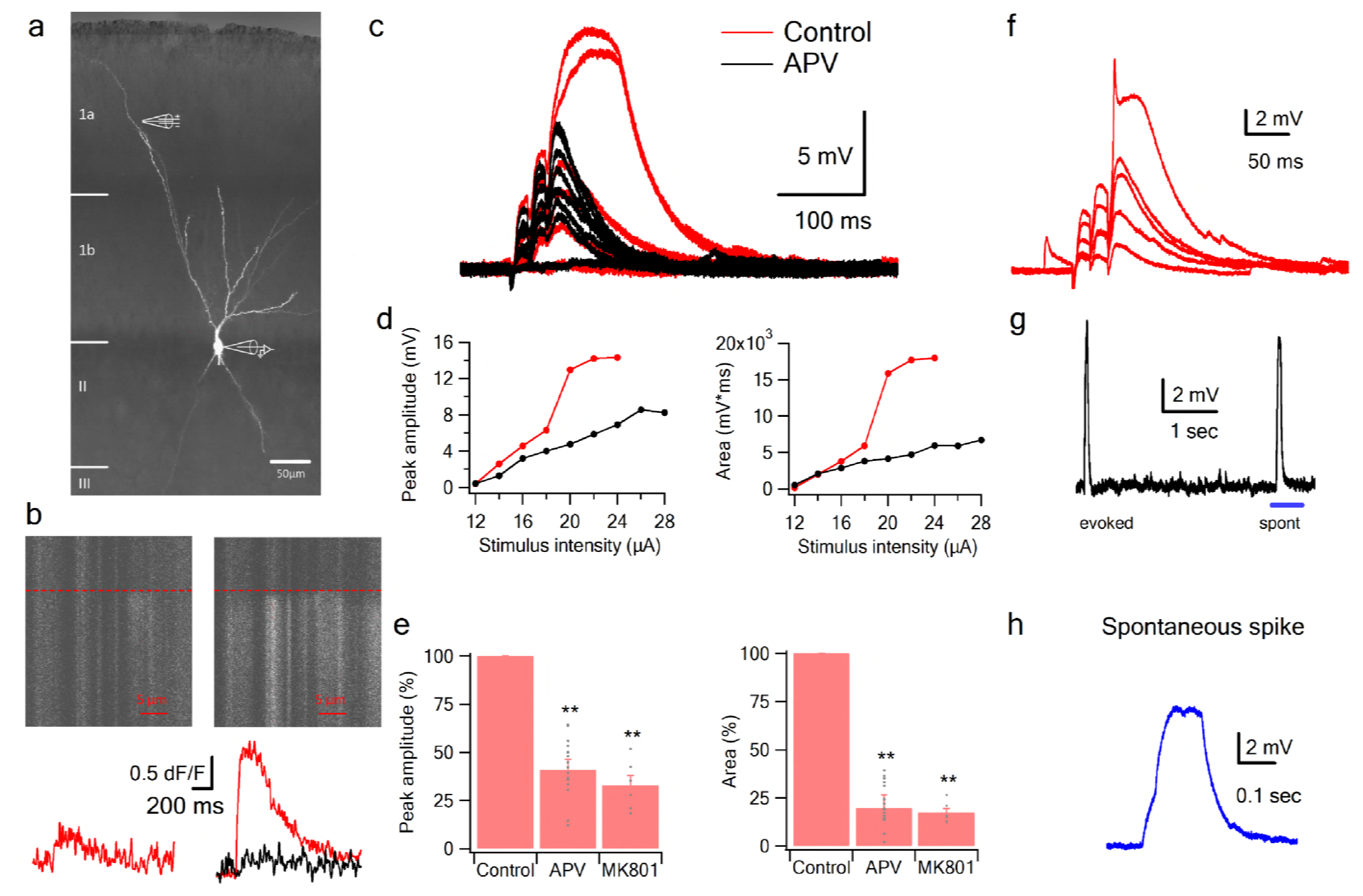
NMDA spikes evoked by LOT stimulation in PCx pyramidal neurons. a. A pyramidal neuron from PCx was loaded with the calcium sensitive dye OGB-6F (200 µm) and CF633 (200 µm) via the patch recording electrode. A focal double-barreled synaptic stimulating theta electrode was placed distally within the LOT innervation zone (280 µm from soma). b. Line scan crossing the dendrite close to the stimulating electrode, showing calcium transients for a subthreshold EPSP (left) and for a dendritic spike at the same site (right). Dashed line denotes the time when the stimulus was delivered. Bottom traces show calcium transients in Control (red) and after addition of APV (black) for a subthreshold EPSP (left) and a dendritic spike (right). c. Voltage responses evoked by gradually increasing synaptic stimulation consisting of a burst of 3 pulses at 50 Hz. With gradually increasing stimulus intensity, an all-or-none response was evoked in control solution (red), which was blocked with the addition of the NMDAR blocker APV (50 µM, black). No biccuculine was added in this experiment. d. Voltage response peak and area plotted as a function of stimulus intensity for the cell shown in A, showing a sigmoidal curve in the Control condition (red) and a linear curve with APV (black). e. Summary plot mean (±SEM) for spike peak amplitude and area in Control (n=48) and after APV application (n=17) or intracellular MK801 (n=6). The tip of the electrode was filled with 1µL of control solution and back-filled with solution containing MK801(1mM). f. Example of a combined NMDA spike and fast spikelet probably representing a local sodium event. g. Example of a spontaneous spike recorded in succession to the synaptically evoked spike denoted by the blue bar is shown with higher time resolution in h. **p<0.01 for comparison with control. Comparison between APV and MK801 did not reach statistical significance.

The average spike threshold evoked by LOT stimulation and recorded at the soma was 14.1±1.6 mV and the dendritic spike amplitude and area under the voltage curve (hereafter “area”) measured at the soma was 27.1 ± 2.4 mV and 3817.8 ± 396.9 mV*ms respectively (mean +/− SEM; n=48 cells; stimulation location 276.78 ± 11.45 µm from soma).

In line with the uncaging data, APV (50 µM) blocked the initiation of dendritic spikes by LOT inputs and linearized stimulus-response curves (Figure 2c-e). At just-suprathreshold stimulus intensity, APV decreased the response peak and area by 56.3 ± 3.8 % and 88.5 ± 2.6 % respectively (n=17). Also similar to uncaging, in 22% of neurons we recorded a fast initial spike component (Figure 2f; amplitude 13.322 ± 0.516 mV, threshold 9.71 ± 0.79 mV, n=8; 217.14 ± 29 µm from soma).

The average resting membrane potential was relatively hyperpolarized (−80.1 ± 1.43 mV), thus in many cases the NMDA spike stayed subthreshold to somatic firing however in other cases we could abserve firing as a result of the NMDA spike.

Occasionally we observed spontaneous spike-like events, which resembled synaptically evoked spikes in shape, including a clear inflection at spike initiation (Figure 2g-h). This indicates that the basic circuitry of the piriform cortex can support the initiation of such spikes. The average spontaneous spike amplitude and area were 17.78±2.09 mV and 9941±2640 mV*ms respectively (n=22 spikes).

To further study the role of NMDARs in synaptically evoked spikes, we blocked NMDARs intracellularly. We and others have previously shown that intracellular MK801 can block NMDARs from the inside thus can serve as a powerful tool to dissect regenerative postsynaptic amplification effects of NMDARs from network recurrent effects (Lavzin et al. 2012). Addition of MK801 to the patch pipette solution completely blocked dendritic spikes (Figure 2e). The spike amplitude and area at just-suprathreshold stimulation intensity was reduced to 32.7±5.3% and 17.2 ± 2.2% respectively (n=6 cells; 40 min after patch breakthrough).

Dendritic calcium imaging revealed that NMDA spikes were accompanied by large calcium transients around the activated dendritic site (Figure 3). Using OGB-6F we observed a maximal calcium transient at the activated dendritic location which fell off steeply in both proximal and distal directions relative to the activated sites (Figure 3b-e). These data are consistent with NMDA spike-evoked calcium profiles seen in basal dendrites of layer 5 pyramidal neurons, and indicate the initiation of a local, non-actively propagated spike ^28,32^.

**Figure 3.**
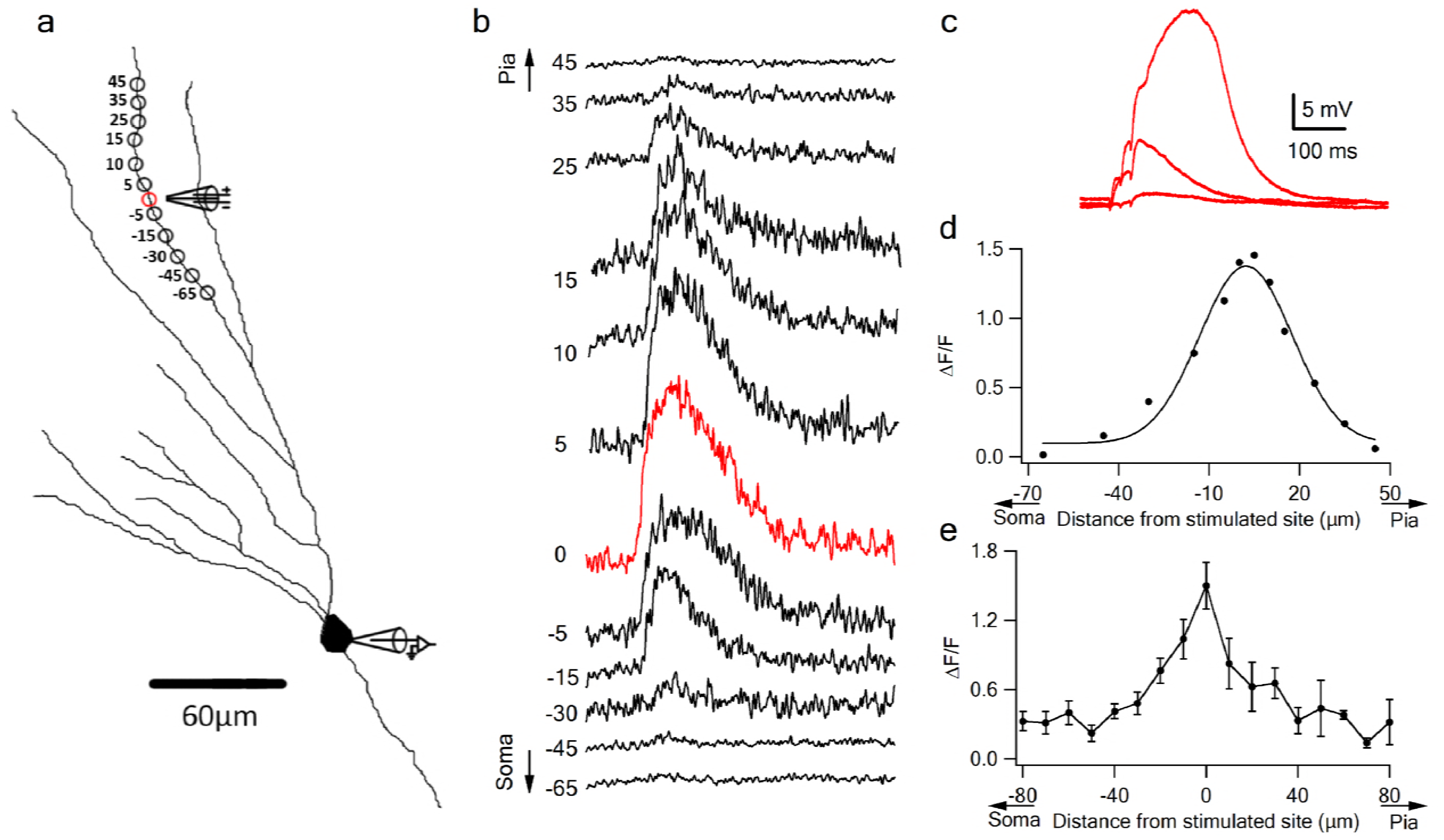
Local calcium transients evoked by dendritic NMDA spikes. a. Fluorescence reconstruction of a layer IIB pyramidal neuron, showing stimulation electrode (288 µm from soma) and the sites of calcium imaging (red circle denotes the location of synaptic stimulation). b. Calcium profile along the stimulated apical dendrite. Calcium transients are expressed as ΔF/F shown for different segments around the stimulated site (as illustrated in A). “0” denotes stimulation location, while distances (in µm) from stimulation site towards pia is indicated as +ve and towards soma as –ve. c. Example of subthreshold EPSPs and an NMDA spike evoked at this recording site. d. Calcium profile (peak ΔF/F) fitted by a Gaussian curve plotted as a function of distance of spike location (for the experiment shown in a-b). e. Summary plot of mean change in calcium transient (ΔF/F ± SEM) evoked by an NMDA spike, as a function of the distance from the center of a stimulated segment (0) averaged in 10 µm segments, from proximal (-ve values) to distal (+ve locations). All cells contained OGB-6F dye, perfused through the patch pipette (n=5).

Together with the uncaging data, these results indicate that LOT inputs are capable of generating NMDA spikes in distal pyramidal neuron dendrites, and these spikes strongly amplify peak and time-averaged somatic EPSP responses compared to just-subthreshold (to NMDA spike) responses (218.4±16 % and 313.5±34.1% for peak amplitude and area respectively).

### NMDA spikes can be generated throughout the apical tree, and by both LOT and IC inputs

In addition to bulb inputs conveyed by the LOT in layer 1a, pyramidal neurons in PCx receive a much larger number of inputs from IC axons in the deeper layers ^36-38^. Typically, an odor response in the piriform cortex is composed of feedforward LOT activity followed by recurrent IC activity, and it is thought that odor responses in PCx are strongly shaped by this recurrent input ^18,39,40^. To develop a more comprehensive picture of dendritic integration in pyramidal neurons of PCx, it is therefore important to determine whether IC inputs can also trigger NMDA spikes in pyramidal neuron dendrites.

Using glutamate uncaging, we first tested whether NMDA spikes could be initiated at progressively more proximal sites along the apical dendrites of PCx pyramidal neurons. We found that spikes could not only be generated throughout the LOT-recipient zone in layer 1a, but also in layers 1b and 2 where pyramidal neurons primarily receive intracortical inputs (Figure 4). In keeping with previous reports (Major et al. 2008), the amplitude of NMDA spikes recorded at the soma increased significantly (from 6.4 ± 0.7 mV to 25.9 ± 3.1 mV) as the uncaging site moved from distal (318.6 ± 7.9 µm) to proximal (99 ± 14.6 µm) dendritic locations (Figure 4c; p=0.00014). Thus, the apical dendrites of PCx pyramidal neurons are capable of generating spikes throughout the layers that receive LOT and IC inputs.

**Figure 4.**
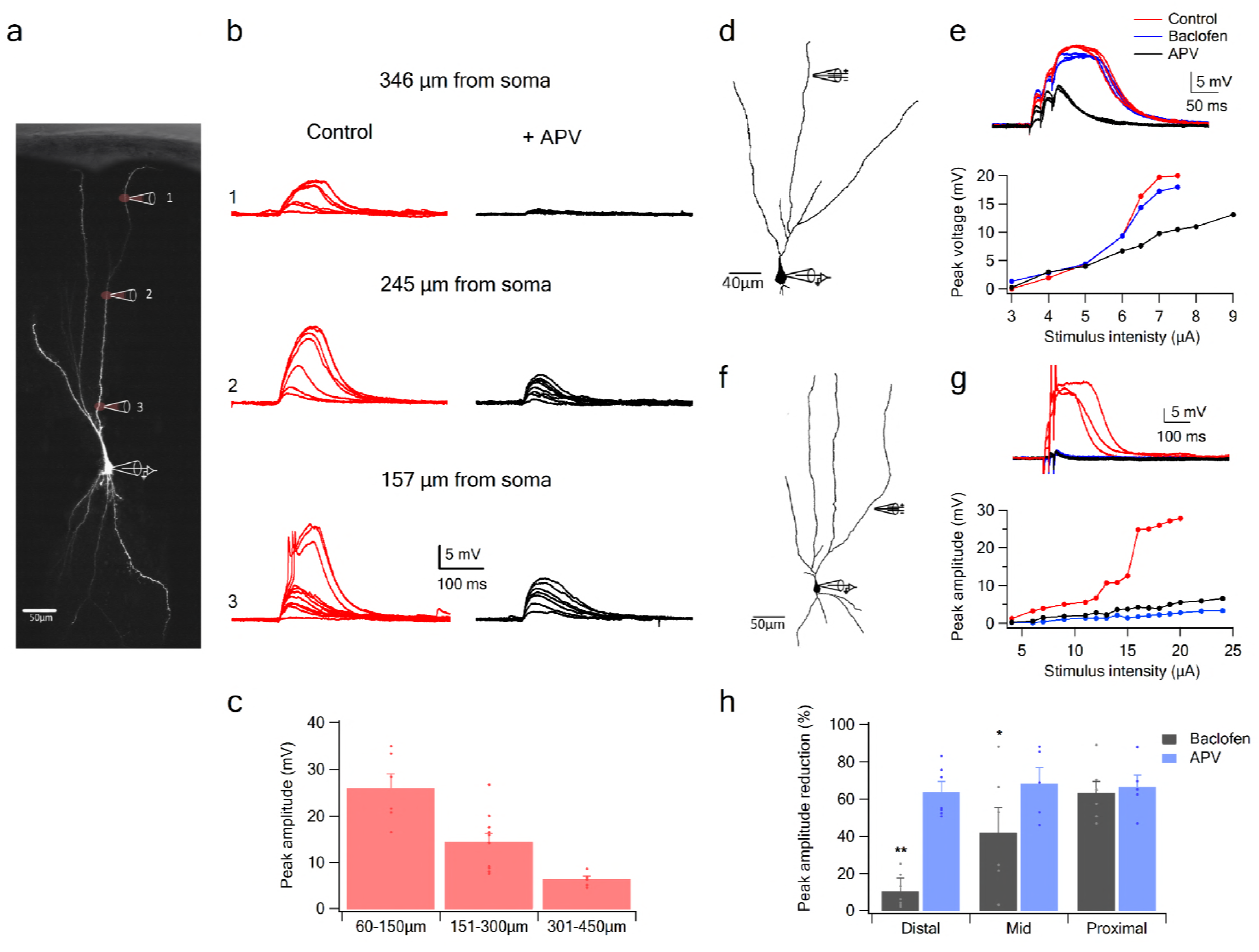
NMDA spike initiation with LOT and IC inputs. a. Pyramidal neuron was loaded with the fluorescent dye CF-633 (200 µm) via the somatic patch recording electrode. Glutamate (MNI-glutamate) was uncaged at three sites as indicated by red circles (346 µM, 245 µM and 157 µm from soma). b. Somatic voltage responses evoked by increasing laser intensity at the dendritic locations indicated in A, in Control (left, red) and with the blocker APV (right, black). c. Summary plot of dendritic spike peak amplitudes as recorded at the soma, as a function of distance from the soma (n=11). d. Reconstruction of a pyramidal neuron showing a focal stimulation electrode at a distal LOT receiving zone (244 µm from soma). e. NMDA spike in Control (red), after sequential addition of Baclofen (100 µM; blue) and APV (50 µM; black). Bottom, plot of peak voltage response as a function of stimulus intensity for Control (red), in the presence of baclofen (blue) and sequential addition of APV (black). f. Reconstruction of a pyramidal neuron showing focal stimulating electrode at a proximal IC dendritic receiving zone (148 µm from soma). g. NMDA spike in Control (red), after sequential addition of Baclofen (100 µM; blue) and APV (50 µM; black). Bottom, plot of peak voltage response as a function of stimulus intensity for Control (red), in the presence of baclofen (blue) and sequential addition of APV (black). h. Summary plot of mean percent reduction in voltage amplitude measured at NMDA spike threshold in control conditions (mean % reduction peak voltage ± SEM) in the presence of baclofen and APV, for distal (n=7), middle (n=6) and proximal (n=6) spike locations. * p<0.05; ** p<0.01

To examine with greater specificity the relative contribution of LOT versus IC inputs to NMDA spike initiation at different distances from the soma, we used the GABA-B agonist baclofen (100 µM), which was previously shown to selectively silence intracortical inputs ^15,35^. When dendritic spikes were initiated at distal dendritic locations using synaptic stimulation (254.85 ± 14.95 µm from soma), addition of baclofen only slightly altered spike amplitude (Figure 4d-e; spike amplitude was reduced by 10.4± 7.3 %, p=0.02 and spike threshold increased by 10.1 ± 12.6 %, p=0.004, n=6). However, at mid and proximal apical dendritic regions, baclofen exerted significant effects on spike initiation and voltage amplitude (Figure 4f-h). At mid dendritic locations (197.3 ± 12 µm), local spikes were evidently triggered by a mixture of LOT and intracortical inputs, since upon baclofen application, response amplitude was reduced (peak voltage response was reduced by 42.0 ± 13.3 %, compared to control; n=6). At more proximal locations (161.6 ± 9.6 µm) spike initiation was almost completely dependent on intracortical inputs: when baclofen was present, we were unable to initiate local spikes at all, and the voltage response was significantly reduced (Figure 4f-g; peak response was reduced by 63.2 ± 6.2 %, compared to control; n=6). At all dendritic locations, addition of APV (50 µm) completely abolished spike initiation (Figure 4d-h). However, at proximal locations APV did not significantly change the response amplitude recorded in the precence of baclofen (Figure 4g-h; reduction of 8 ± 10.6 %; p=0.44; n=5). Thus, mainly the regenerative part of the spike was suppressed by the intracortical blockade by baclofen, leaving only the underlying EPSP.

Taken together these results indicate that pyramidal neurons in PCx are capable of NMDA spike generation throughout their apical arbors; that both LOT and IC inputs can generate NMDA spikes within their respective layers; and that these synaptically evoked dendritic spikes show the commonly observed increase in amplitude as the initiation site moves closer to the soma ^28^.

### Combination selectivity and compartmentalization of pyramidal neuron dendrites

Having a “discontinuous” receptive field means an olfactory pyramidal neuron must respond selectively to multiple different combinations of LOT inputs (e.g. AB and CD), but not to the individual inputs (A, B, C, or D), or to re-combinations of the same inputs (AC or BD). We tested a pyramidal neuron’s capacity for responding selectively to multiple distinct LOT input combinations, in two stages.

First, we verified that the NMDA spike thresholding nonlinearity could provide a mechanism for enforcing combination selectivity *within* a dendrite. For example, a dendrite with a threshold of 2 could respond to a combination of inputs 1 and 2, but not to the individual components 1 or 2. To test this, we used stimulating electrodes to activate two LOT inputs separately and in combination on a single dendrite (3 EPSPs at 50 Hz; 354.4 ± 17.47 µm from soma; average interelectrode distance of 35.69 ± 1µm). Input 1 was activated over a full range of intensities, up through NMDA spike initiation. After generating input 1’s baseline input-output curve, the same stimulus sequence was repeated in the presence of input 2, which provided a constant “bias” input (average EPSP bias amplitude at the soma was 3.81 ± 0.24 mV; n=26). Coactivation of the two inputs resulted in a strong nonlinear interaction, where input 2 significantly lowered the threshold for local spike generation by input 1 (by 46.55 ± 1.8% for peak voltage; Figure 5a-d) without changing the response peak amplitude. The pronounced left shift of the input-output curve caused by input 2 is a fundamentally nonlinear interaction resembling the function *sigmoid*(x_1_ + x_2_, θ), where x_1_ and x_2_ represent the magnitudes of the two inputs, and θ represents the threshold. Had the interaction been linear, the effect of input 2 would have been to lift input 1’s entire input-ouptut curve vertically by an amount equal to the bias voltage, with no change in threshold, giving the *sigmoid* (x_1_, θ) + x_2_. Given the form of the within-branch nonlinearity, we conclude that with an appropriate setting of the NMDA spike threshold, the distal dendrites of olfactory pyramidal neurons are well suited to enforce LOT combination-selectivity.

**Figure 5.**
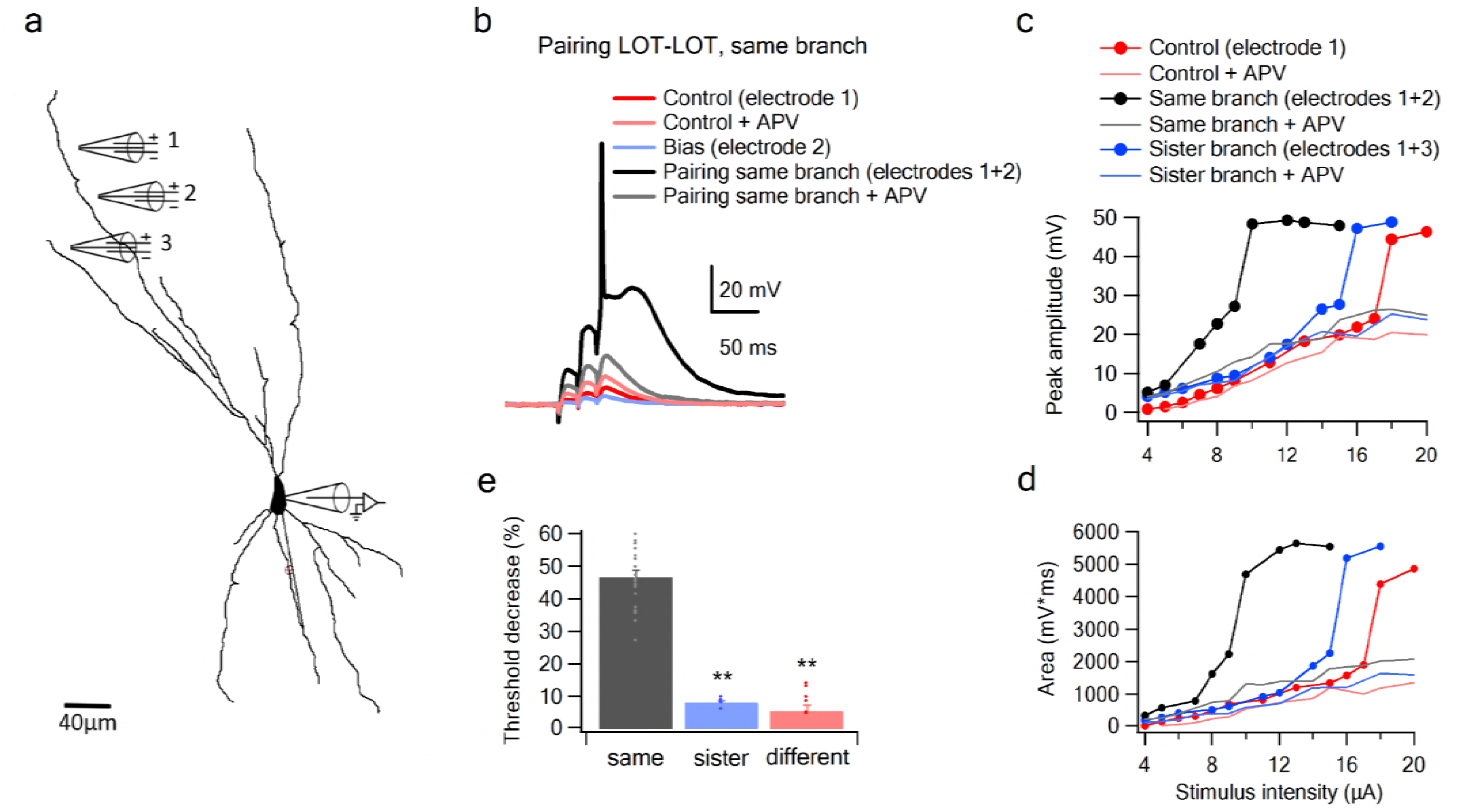
Summation of LOT inputs on same, sister and different dendritic branches. a. Reconstruction of a layer IIB pyramidal neuron filled with the fluorescent dye CF-633 (200 µM) with stimulating electrodes positioned within an LOT-receiving zone in same dendritic branch (electrodes 1, 2) and sister branch (electrode 3). b. Voltage responses to pairing LOT inputs in same branch (electrodes 1+2) in Control (black) and with APV (grey). Example responses to electrode 1 separately (red), electrode 2 (blue, bias voltage), and electrode 1 with APV (light red) are shown. c. Stimulus response curves (peak voltage) for paired LOT inputs within the same branch (electrodes 1+2; inter-electrode distance 38 µm) and pairing with a sister branch (electrode 1+3). d. Same as in C, for area of the responses. e. Summary plot of percent decrease of spike threshold in the paired condition relative to Control for LOT-LOT inputs in same branch (grey) sister branch (blue) and different branch (red). ** p<0.01 for comparison with same branch. Comparison of sister and different branches did not reach statistical significance (p=0.24). Sister branches were defined as branching from same dendrite; Different branches were defined as branching from two separate branches.

The second stage question is whether a co-activation of two inputs of comparable magnitude to those used in the experiments above, but split *between* dendrites, are less effective at driving the cell than the within-branch combination. We found in favor of this hypothesis that the nonlinear interaction between LOT inputs delivered to two different branches, even when those branches were “sisters” (Figure 5c-e), was much weaker than the within-branch interaction, as evidenced by the much smaller change in spike threshold caused by the bias input (7.98 ± 0.75%, n=5 threshold change for the sister-branch Figure 5c-e). The threshold coupling between branches was weaker still when the bias was delivered to a different branch that was more remote than a sister (6.28 ± 1.88%, n=6 threshold change for the different-branch, Figure 5e).

To more closely examine the role of NMDA channels in the pairing outcomes, we repeated the pairing experiments in the presence of APV (Figure 5c-d). In these cases, the bias added a roughly constant value to the unpaired voltage response (Control + APV) all along the curve, confirming that without NMDA channels, pyramidal neuron dendrites revert to roughly linear summation.

Together these results support a model of olfactory coding in PCx in which (1) NMDA spike generation in the distal apical dendrites of pyramidal neurons provides the superlinearity needed to enforce selectivity for specific combinations of LOT inputs, while the compartmentalization of voltage signals in the apical tree allows for different LOT combinations to be mapped onto different apical dendrites with relatively little crosstalk between them.

### Nonlinear summation of LOT and IC inputs

As odor responses in pyramidal neurons of the PCx are driven first by direct LOT activity and subsequently shaped by recurrent IC inputs ^4,9,39,40^, it is critical to understand how LOT and IC inputs summate, whether linearly or nonlinearly, and if nonlinearly, with what type of nonlinear interaction.

Since LOT and IC inputs are segregated along the proximal-distal axis of a pyramidal neuron’s apical dendritic tree, we used focal synaptic stimulations to examine the interaction between distal sites representing predominantly LOT inputs, and more proximal sites representing predominantly IC inputs. In these experiments, one electrode was positioned at a distal site and held fixed, while a second electrode, which again provided the bias input, was moved closer to the soma within the same branch (Figure 6a; average interelectrode distance of 138.46 ± 7.78µm and bias voltage of 3.9±0.27 mV; n=13). We found that summation of LOT and IC inputs was very similar in form to the summation of two inputs confined to the LOT (Figure 6a-e): the IC bias input again led to a substantial threshold reduction for LOT inputs (35.44 ± 2.0%, n=15), with no increase in response magnitude. Indeed, the threshold-lowering effect of the bias input was relatively constant within increasing separation of the two inputs (Figure 6f), indicating that over much of its length, a pyramidal neuron apical dendrite functions as a single, relatively location-insensitive integrative subunit.

**Figure 6.**
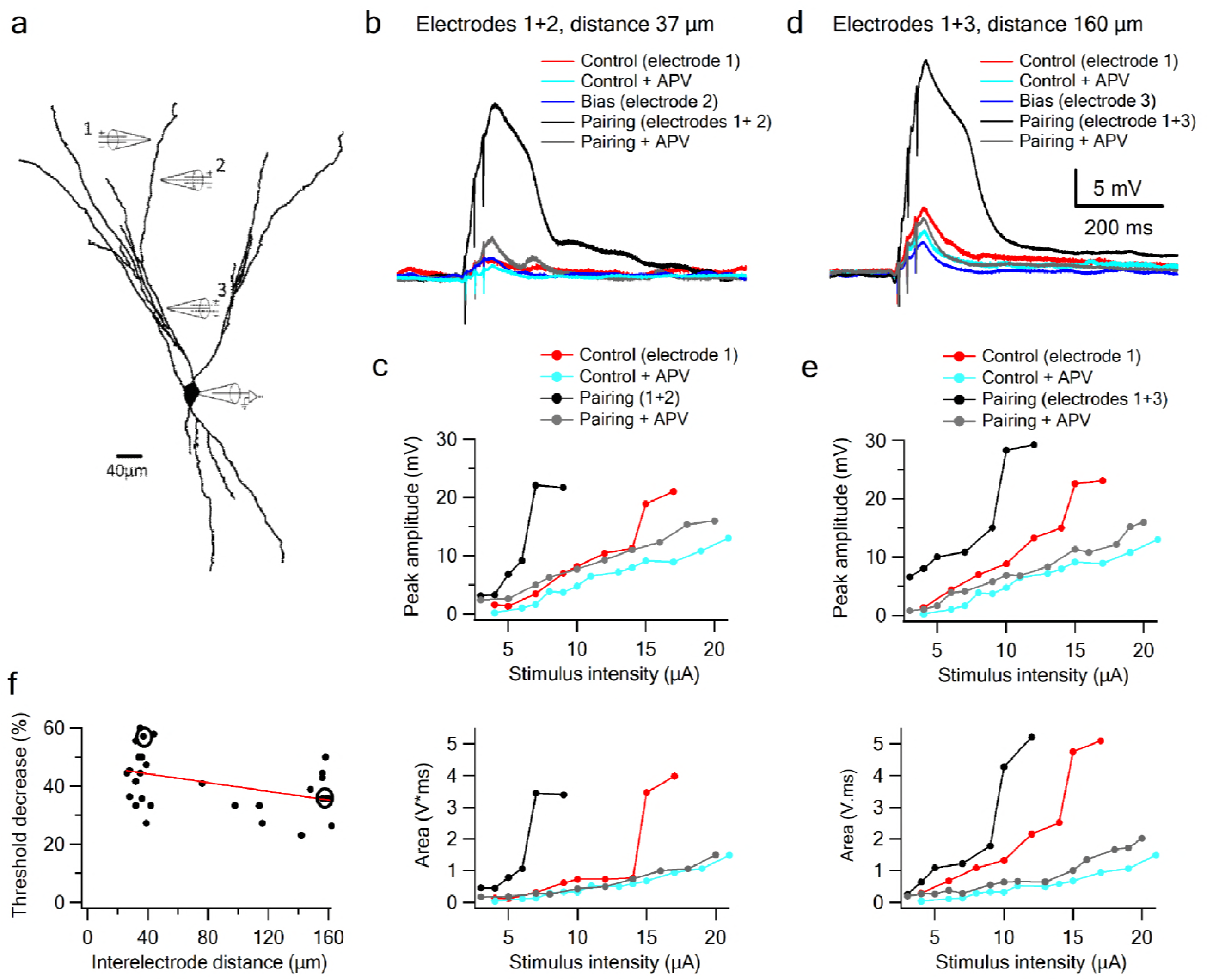
Supralinear summation of LOT and IC inputs on a single dendritic branch. a. Reconstruction of a layer III pyramidal neuron filled with the fluorescent dye CF-633 (200 µM). Three stimulating electrodes where positioned in close proximity to same dendritic branch, with electrodes 1 and 2 within the LOT-receiving zone and electrode 3 within the IC-receiving zone. b. Voltage responses for paired LOT inputs on the same branch (electrodes 1 and 2, inter-electrode distance 37 µm). Example responses are shown for electrode 1 (red), electrode 1 + APV (cyan), electrode 2 (blue, bias voltage), pairing activation of electrodes 1+2 (black), and pairing activation of electrodes 1+2 in the presence of APV. c. Stimulus response curves for LOT pairing on the same branch (top voltage; bottom area). d. Same as b, for pairing LOT and IC inputs on the same branch (electrodes 1 and 3, inter-electrode distance 160 µm). e. Stimulus response curves for LOT and IC pairing on same branch (top voltage; bottom area). f. Summary plot of percent decrease of spike threshold (paired condition relative to control) as a function of inter-electrode distance in same branch. Circles show the examples shown in the figure. Slope= −0.075 ± 0.03 mV/µm

In contrast to the weak dependence on location, the nonlinear interaction of LOT and IC inputs depended completely on NMDA regenerativity: blocking NMDARs linearized the input-output curves, destroying their sigmoidal form, and eliminating the basis for a superlinear within–branch interactions (Figure 6, grey curves).

To complete the picture, we examined the interaction between a distal LOT input and a proximal IC input either on sister or different branches (Figure 7). When the bias was provided by an IC input on a sister branch, the nonlinear interaction was evident, though significantly weaker compared to that seen with a same-branch IC bias (Figure 7a-b, e). Threshold reduction was 23.38 ± 2.34% (n=8) for IC locations on sister branches. The interaction was weaker still when the IC bias was on a more distantly related branch, 18.4 ± 2.88% (n=7), approaching the minimal nonlinear interaction seen with LOT-LOT different-branch pairing (Figure 7c-e).

**Figure 7.**
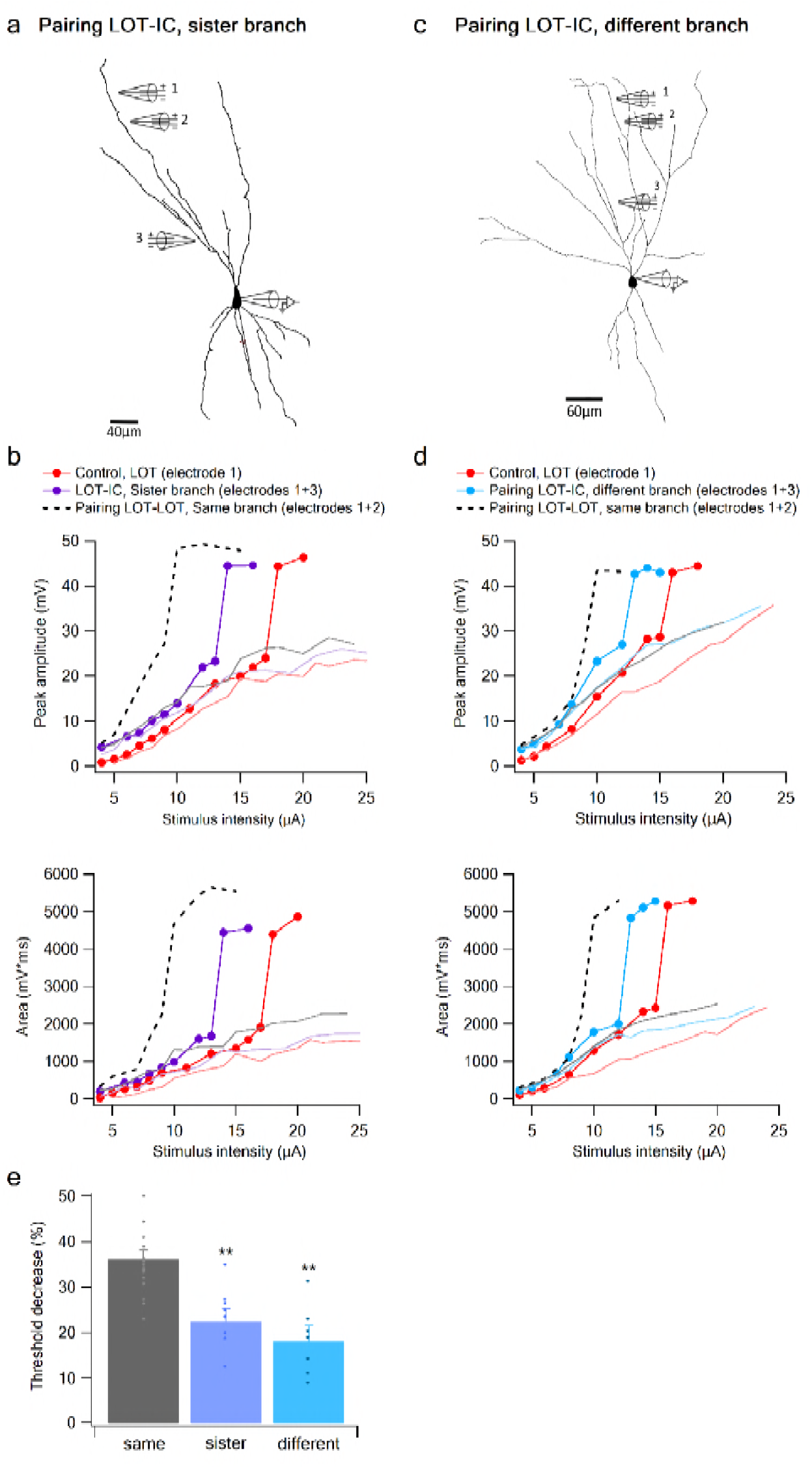
Summation of LOT and IC inputs on sister and different dendritic branches. a. Reconstruction of a layer IIB pyramidal neuron (same as in Figure 6) filled with the fluorescent dye CF-633 (200 µM) with electrodes positioned within LOT regions of the same branch (electrodes 1,2), or at IC-receiving dendritic regions at sister branch (electrode 3). b. Stimulus response curves for a single electrode (1, red), paired LOT-IC responses located on sister branches (1+3, purple), shown for peak amplitude (top) and area (bottom). Solid lines show same responses in the presence of APV. For comparison the paired LOT-LOT curve on the same branch (dotted black) is shown. c. Reconstruction of a layer IIB pyramidal neuron filled with the fluorescent dye CF-633 (200 µM) with electrodes positioned within the LOT-receiving region of the same branch (electrodes 1,2), or at IC-receiving dendritic region at different branch (electrode 3). d. Same as in B but for LOT and IC inputs activated on different branches. e. Summary plot of percent decrease of spike threshold in paired condition relative to control for LOT-IC inputs to the same branch (grey), sister branch (blue), and different branch (teal). ** p<0.01

In summary, summation between LOT-IC inputs within a single apical dendrite is also nonlinear, with only a slight weakening of the nonlinear interaction as the inputs are increasingly separated (i.e. compared to more closely spaced LOT-LOT pairs). This indicates that apical dendrites of pyramidal neurons in PCx, function as relatively simple separate integrative subunits that apply a sigmoidal nonlinearity to their summed inputs with relatively little dependence on location. On the other hand, we observed a much weaker nonlinear interaction between inputs to different dendrites, indicating pyramidal dendrites enjoy a significant degree of dendritic compartmentalization.

### Modeling

To cross check our experimental findings, we developed a compartmental model of a reconstructed PCx pyramidal neuron, and recorded its responses to stimulus configurations similar to those used in our experiments. We first established that the model cell can generate NMDA spikes in response to concentrated synaptic excitation at any location along a pyramidal neuron’s apical dendrite, and that both the threshold for spike initiation and the spike amplitude measured at the soma increase as the site of spike initiation moves closer to the soma (Supplementary Figure S1). We next replicated the same/sister/different branch pairing experiments shown in Figures 5-7. We found a close correspondence between the experimental and modeling results, wherein an LOT bias input activated on the same dendrite produced a much larger threshold-lowering effect than a bias input of the same size (measured at the soma) delivered to the LOT region of a sister or different branch (Supplementary Figure S2). Thus, the model supports our experimental finding that nonlinear synaptic summation of LOT inputs to distal apical dendrites is strongly compartmentalized, with individual apical dendrites acting as well-separated integrative subunits.

We also found close correspondence to the experimental data for interactions between LOT and IC inputs on same, sister and different branches (Supplementary Figure S2). The threshold lowering power of an IC bias input on the same branch was somewhat reduced compared to a bias input activated within the LOT itself, consistent with a mild distance-dependent attenuation of synaptic interactions on pyramidal neurons on same apical dendrites. In addition, the model replicated the experimental finding that IC bias inputs on sister and different branches had a significantly weaker effect than an IC bias on the same branch (compare Supplementary Figure S2g to Figure 8e), though the degree of compartmentalization of IC inputs was, as in the experiments, and as expected from passive cable theory, less pronounced than the compartmentalization of LOT inputs. Thus, the model supports our experimental findings that IC inputs interact nonlinearly with LOT inputs, but compared to highly compartmentalized interactions between LOT sites on different branches, the effects of IC inputs are less well compartmentalized.

**Figure 8.**
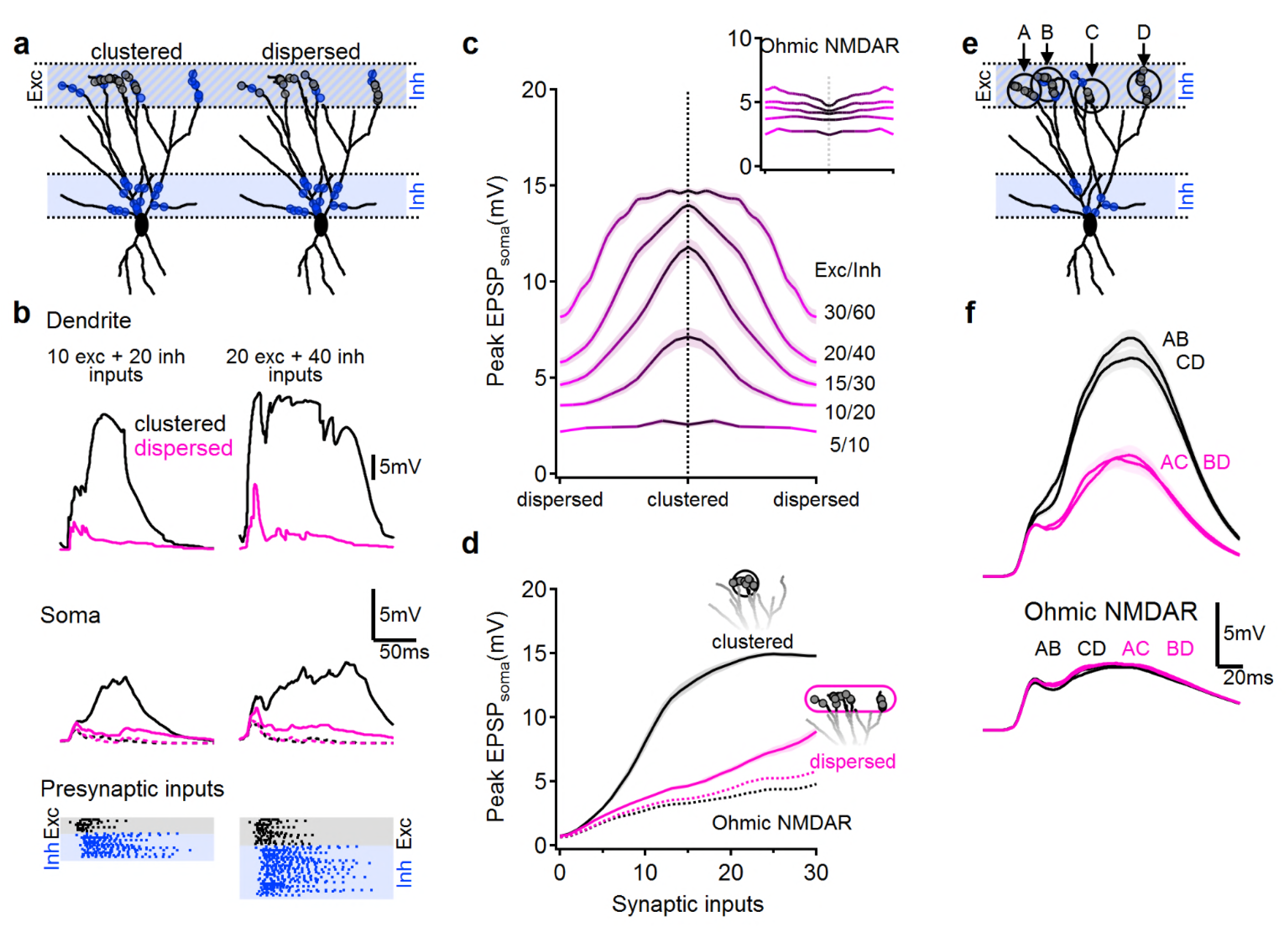
NMDA spikes can produce combination-sensitive dendritic receptive fields: modeling results. a. Example distribution of 20 glutamatergic inputs (grey circles) and 40 GABAergic inputs (blue) on the reconstructed cell (for clarity, some dendrites are not shown). Left, clustered distribution, where excitatory inputs are concentrated on a single postsynaptic branch. Right, dispersed excitatory distribution over all distal dendrites. b. Example postsynaptic responses to stimulation with 10/20 (left) and 20/40 (right) presynaptic excitatory/inhibitory inputs. Top, voltages recorded from the stimulated dendrite. Middle, somatic EPSPs. Dotted traces – stimulation in the Ohmic NMDARs. Bottom, the temporal presynaptic activation pattern, simulated to mimic typical odor responses of mitral cells *in-vivo*. The presynaptic firing trains were identical between clustered and dispersed distributions. c. Peak somatic EPSPs as a function of input clustering for different number of synaptic inputs. d. The simulated peak EPSP amplitude recorded at the soma as a function of the number of presynaptic inputs. Blockage of NMDA spikes with Ohmic NMDA channels abolished the preference for clustered excitatory drive (n=100 repetitions for each stimulation intensity). e-f. Dendritic odorant selectivity with NMDA spikes. e. Distribution of excitatory inputs (black) from four glomeruli (A-D, 10 inputs each) and odor-unselective inhibitory background (40 synapses, blue). f. Example somatic voltage profiles following activation of different combinations of LOT inputs. Top, stimulation of glomerular inputs that terminate on the same primary dendrite (black) promoted stronger postsynaptic depolarization compared to a dispersed input combination (magenta). Bottom, combination selectivity disappeared when NMDA spikes were blocked with Ohmic NMDA channels (n=100 repeats for each condition). Shaded areas-SEM. See also Supplemental Figure S1 and S2.

Finally, we used the model to verify that under in vivo-like conditions, the combined effects of NMDA spikes and dendritic compartmentalization can produce the “discontinuous” receptive fields typical of pyramidal neurons in PCx. In particular, we predicted that combination selectivity would be observed for glomerular activation patterns that resulted in clustered excitation on apical dendrites, since this would tend to activate NMDA spikes and powerfully drive the cell, whereas input combinations that activated apical dendrites diffusely would fail to trigger NMDA spikes and therefore drive the pyramidal neuron only weakly.

To test this idea, we distributed LOT inputs in the distal apical tree in either a clustered or dispersed fashion, along with inhibitory inputs targeting both the distal apical dendrites (representing feedforward inhibition), as well as the perisomatic region (representing feedback inhibition) (Figure 8a). We activated LOT inputs with in vivo-like firing patterns designed to mimic mitral cell responses to odors (Figure 8b, bottom;^9,18,41,42^). The timing and kinetics of feedforward and feedback inhibitory inputs were likewise designed to mimic in-vivo like activation as described in the literature (Figure 8b) ^18,43-47^.

We compared clustered activation of LOT inputs (Figure 8, black) to the same number of synaptic inputs dispersed randomly over the entire LOT-receiving area of the dendritic tree (Figure 8, pink). As predicted, we found that that clustered LOT inputs reliably evoked NMDA spikes, powerfully amplifying postsynaptic signal compared to Ohmic (voltage-independent) NMDAR (Figure 8c-d). In contrast, dispersed LOT activation typically failed to overcome the local spike threshold on any dendrite, due both to the lack of concentrated excitation and the presence of inhibition, and therefore resulted in significantly smaller local dendritic and somatic voltage responses (Figure 8b-d, pink). The difference between clustered and dispersed synaptic distributions was evident over a large input range (Figure 8c-d), leading to preferential amplification of input combinations that target individual postsynaptic branches (Figure 8c). This effect was entirely dependent on regenerative NMDAR currents: stimulation with Ohmic NMDARs, eliminated the strong dependence of postsynaptic responses on the spatial distribution of synaptic inputs (Figure 8c, inset, 8d, dotted).

## Discussion

How pyramidal neurons in piriform cortex integrate their bulb inputs to generate olfactory percepts has been an unsettled question. To address this, we studied the integrative properties of pyramidal neuron dendrites in PCx using glutamate uncaging, focal synaptic stimulation, and compartmental models. Our primary aim was to determine whether local spike generation in the dendrites of PCx pyramidal neurons could serve the dual purposes of amplifying pyramidal neuron responses to LOT inputs impinging on their distal dendrites, as well as provide a nonlinear binding mechanism that could underlie a pyramidal neuron’s selectivity for multiple distinct odorant combinations.

We found that pyramidal neurons in PCx can generate local spikes throughout their apical dendrites, and that a spike in a dendritic branch can powerfully depolarize the soma (i.e. producing up to 40-50 mV depolarizations depending on distance and cell size). Our results are consistent with two key properties of local spikes seen in thin (basal, apical oblique, and tuft) dendrites of pyramidal neurons in other cortical areas. First, the majority of spikes we observed in PCx pyramidal neurons are mediated primarily by NMDAR channels similar to thin dendrites in other pyramidal neurons ^20,21,24,27,29,32-34^. In addition, in a subset of cases we observed fast spikes reminiscent of sodium spikelets observed in thin tuft and basal dendrites of pyramidal neurons in other cortical areas ^21,23,28,33^. Second, the NMDA spike amplitude (measured at the soma) increase progressively as the site of spike initiation moves closer to the soma ^28,48^.

Beyond the properties of NMDA spikes per se, we found that the logic of synaptic integration in pyramidal neuron dendrites in PCx, particularly the pronounced difference between within-branch and between-branch summation, is also consistent with that seen in other types of pyramidal neurons ^18,19,49^. In particular, synaptic inputs to a PCx pyramidal neurons are processed via a 2-layer computation. First, LOT inputs are combined within individual dendrites as roughly a weighted sum (i.e. linearly) up to the local spike threshold (Figure 2d), so that an individual dendrite behaves comparably to a “neuron”in a conventional artificial neural network. A telltale sign of this type of linear-nonlinear (LN) input-output function *within* a dendrite is the nearly pure left-shifting of a “control” input’s sigmoidal input-output curve caused by a constant “bias” input activated on the same branch – an effect seen in both our experimental data and simulation results (Figure 5, and Supplementary Figure S2). In the second layer of processing, the outflows from the separately thresholded dendritic “subunits” are combined linearly at the soma as a prelude to output spike generation. A telltale sign of linear summation *between* dendrites is the uniform lifting of one branch’s input-output curve (especially over its subthreshold range where saturation effects are minimal) by the constant somatic bias voltage generated by another dendrite. This is best seen in the peak amplitude curves in Figure 7b and d, and in the model input pairing figure (Supplementary Figure S2). A second sign of (relatively) independent functioning of different dendrites is the much smaller threshold shift in a dendrite’s i/o curve seen when a bias input is applied to a different branch, where the least nonlinear crosstalk occurs between LOT-receiving zones in “cousin” branches, and only slightly more between directly adjoining “sister” branches (Figures 5e, 7e, and Supplementary Figure S2g; see also ^50^).

The 2-layer architecture of a pyramidal neuron in PCx allows it to respond selectively to specific high-order odorant combinations – those whose LOT activity patterns deliver concentrated (suprathreshold) excitation to at least one apical dendrite – without responding to the vast majority of LOT patterns that produce more diffuse, and therefore subthreshold, excitation to multiple branches within the dendritic arbor. This ability to respond to multiple distinct high-order combinations, without responding to re-combinations of the same odor components, may account for this cell type’s hallmark physiological property, namely its “discontinuous” receptive field ^13,16-18^. This same type of scenario, in which a neuron computes a disjunction over a set of nonlinear “features” mapped onto different dendrites has been previously proposed to underlie the pooling of multiple simple cell-like subunits within a complex cell’s receptive field in V1 ^49,51^; the pooling of higher-order feature conjunctions in a memory circuit ^19,52-54^, and as a means to multiplex one of several neural pathways through to a cell’s output, as might occur in the context of a decision task ^55^.

### Mismatch to the currently accepted view of olfactory pyramidal neurons

As previously discussed, our results showing that pyramidal neurons in PCx have a 2-layer summation logic arising from (1) NMDA regenerativity and (2) a compartmentalized dendritic tree, are inconsistent with a previous study in olfactory cortex which reported little sign of regenerative NMDA or sodium currents in pyramidal neuron dendrites, and concluded that pyramidal neurons in PCx integrate their inputs essentially linearly ^26^. The reasons for the discrepancy between the Bathellier et al. study and ours are unknown, but presumably stem from methodological differences. One difference in the experimental conditions was the slicing procedure: We used coronal slices whereas Bathellier et al. used parasagittal slices. This may have led to differences in the preservation of LOT inputs in our slices, which may have increased the efficacy by which dendrites could be stimulated focally.

### Transformation of the odor code from olfactory bulb to piriform cortex

Anatomical data indicate that (1) axons originating in the olfactory bulb and traveling along the LOT terminate broadly throughout the piriform cortex, targeting a dispersed population of pyramidal neurons, and (2) single pyramidal neurons receive inputs from multiple broadly distributed olfactory glomeruli ^8,10,11^. Lacking any evidence for a patterning of these connections, the anatomical projection from the olfactory bulb to the cortex is generally assumed to be random. In keeping with this assumption, physiological data show that different odors activate a unique set of neurons widely distributed across the PCx, and that individual olfactory pyramidal neurons respond to a small, unpredictable subset of odors ^13^. Thus, whereas the bulb forms a molecular-based code wherein M/T cells in a given glomerulus respond to any odor containing that glomerulus’ associated molecule, pyramidal neurons in PCx fire only to high-order glomerular combinations, but respond to multiple combinations that have little to no chemical overlap with each other.

This transformation from molecule-specific responses in olfactory glomeruli to multi-combination selectivity in pyramidal neurons in PCx requires a nonlinear transformation, and indeed several in-vivo and in-vitro studies have indicated that single pyramidal neurons in PCx do integrate odor information nonlinearly ^9,13,15,18^. In particular, activation of a single glomerulus or LOT fiber generates only a small subthreshold depolarization in most connected PCx pyramidal neurons. On the other hand, when multiple glomeruli or multiple LOT inputs are activated simultaneously, or combinations of odorants are presented to an animal, PCx pyramidal neurons produce strong supralinear responses ^9,13,15,17,56^.

What mechanism underlies the supralinear integration of convergent LOT inputs onto single PCx pyramidal neurons? Apicella et al. ^15^ showed the supralinearity requires neither cooperative interactions in the bulb, nor participation of recurrent IC inputs in PCx. Another potential source of supralinearity would be the cell’s output spiking mechanism: a high threshold for somatic action potential generation could in principle be used to limit a pyramidal neuron’s responses to only those stimuli that activate N (or more) connected LOT axons. However, without dendritic subunitization, the cell should respond to *any* combination of N LOT inputs, destroying the combination selectivity needed to account for a PCx pyramidal neuron’s discontinuous RF. In contrast, our results support the idea that the dendritic thresholding nonlinearity provided primarily by regenerative NMDA currents can mediate the supralinear integration of LOT inputs observed previously both in-vitro and in vivo.

### The softer compartmentalization of IC inputs, and its implications

In our exploration of the nonlinear interactions between driver inputs within the LOT and bias inputs delivered either within the LOT or at the IC-receiving regions of the apical tree, we found that the threshold-lowering effects of IC inputs were less well compartmentalized. The effect can be traced to passive cable theory: IC inputs are closer to the branch points where sister and cousin dendrites connect to each other, so that their effects are felt more widely. In quantitative terms, beginning with a “control” input-output curve generated by an LOT input, we found that the threshold-lowering power of a second LOT input on the same branch was roughly 10 times that of an LOT input delivered to a different branch. In contrast to this strong compartmentalization, the threshold-lowering effect of an IC bias input on the same branch is only twice that of an IC bias delivered to a different branch (Figure 7e). The observation that IC inputs modulate more globally comes with a caveat, however: it was previously shown that the degree of nonlinear crosstalk between dendritic branches tends to be overestimated in subthreshold summation experiments, compared to a cell operating in the firing regime which enhances subunit independence ^50^. This is because the somatic spike-generating mechanism acts as a sort of time-averaged “voltage clamp” ^57^ that suppresses subthreshold voltage communication between dendrites ^50^. In light of this effect, it remains to be determined whether the softer compartmentalization of IC inputs seen in both our experiments and simulations will persist to the same degree under normal operating conditions in the olfactory cortex.

Both the existence of nonlinear interactions between feedforward LOT and recurrent IC inputs to PCx pyramidal neurons, and the (unknown) degree to which IC inputs act locally (i.e., have modulatory effects confined to a single dendrite) vs. globally (affecting some or all dendrites) in vivo, suggest there remains much to learn about the functioning of the recurrent odor recognition network in piriform cortex. Besides carrying feedback from other pyramidal neurons in the area, IC inputs provide contextual information from higher-order cortical regions including the entorhinal cortex, orbitofrontal cortex and amygdala, potentially allowing the assignment of cognitive and emotional value to odors ^1,38^. The nonlinear interaction of LOT and IC inputs mediated by NMDA regenerativity could provide a biophysical mechanism for binding odor with contextual information in piriform cortex. If so, the dendritic subunitization of PCx pyramidal neurons, and the possibility of some locality of IC modulation within a neuron, could allow contextual information to be bound to certain odorant combinations represented by a neuron and not others.

### Possible role of NMDA spikes in dendrite-specific plasticity induction

Backpropagating action potentials in apical dendrites of PCx pyramidal neurons attenuate significantly as they propagate ^26,58^, making it less likely that bAPs contribute to spike timing dependent plasticity of distal LOT synapses. In contrast, given that NMDA currents can produce large localized calcium transients in apical dendrites, confined to within ± 20 µm of the activated site, such spikes could serve as local induction signals for plasticity of LOT synapses. In accordance with this notion, it was recently shown that NMDA spikes contribute to long-term potentiation (LTP) in the dendrites of CA3 neurons ^59^. In the same way, a group of LOT synapses that fire together on the same dendrite in PCx could trigger a local plasticity event that induces LTP of the activated synapses. When the same odor is re-encountered at a later time, and re-activates the group of now potentiated synapses, an even more powerful NMDA-dependent response might be generated, signaling odor recognition (for a discussion of related ideas see ^52,60^). Further work will be needed to determine the ways and conditions in which synaptic plasticity contributes to the learning-related functions of the olfactory cortex.

## Author contribution

“A.K., O. S. and J. S. conducted the experiments and performed the analysis; J. S. conceptualized and designed the experiments and performed analysis and figures; A.P. and B.M. performed the modeling and figures; J.S., B.M., A.P. and E.B. wrote the paper.

## Acknowledgments

We thank Y. Schiller for helpful discussions throughout the project and helpful comments on the manuscript. We thank Irena Reiter for excellent technical assistance and processing the biocytin-filled neurons. This study was supported by Israeli Science Foundation (J.S.), the Rappaport Foundation (J.S.), the Adelis Fund for Brain Research at the Technion and Price funds (J.S.).

---

**Supplementary Figure S1. NMDA spikes in PCx pyramidal neurons: modeling results.**

a. Representative examples of focal synaptic stimulation at the LOT (top) and the IC (bottom) regions. b. AMPAR only stimulation, same locations. Colors code synaptic strength as labeled. c. Peak somatic EPSP as a function of synaptic conductance for the two dendritic locations, LOT (top) IC (bottom). d. Dependence of NMDA spike threshold (red) and NMDA spike amplitude (pink) on the distance of the stimulation site from the soma.

**Supplementary Figure S2. Related to Figure 8. Pairing LOT and IC inputs: modeling results.** a-f. Example paring of a glutamatergic synaptic input on a distal apical branch (control) and a bias input located on the same branch (a, b), sister branch (c, d) or ‘different’ dendrite (i.e. that do not share a primary branch, e, f). Left, top – morphology of the reconstructed cell and stimulation locations. Left, bottom, example paring; red – somatic EPSP following activation of 5 nS control input; blue, EPSP following activation of a 4 nS bias, this bias level was selected to produce ∼5 mV depolarization at the soma and ∼50% reduction in NMDA spike threshold for paring of nearby inputs, similar to experimental conditions. Black, combined activation of bias and control synapses. Dotted line, the expected linear summation of the two individual inputs. Right, peak somatic EPSP amplitudes for progressively increasing stimulation intensities of the control input. Right bold trace – in the absence of bias, lighter traces – simulations in the presence of a bias input; traces differ by the magnitude of the bias intensity (left bold trace, bias of 4 nS). Arrow indicated the stimulation intensity of the control input used to produce the left, bottom plots. g. Reduction in spike threshold between control activation only and pairing of control with 4 nS bias inputs as a function of distance between the two stimulation locations. Data from 6 different control locations in 3 reconstructed cells. Inset, schematic locations of the bias input color coded by same/sister/different dendrites, location of the control input is as in a-f; also depicted here by the large circle.

## Materials and Methods

### Electrophysiology and calcium imaging

Coronal brain slices 300 µm thick from a 28-40 day old Wistar rats (male and femal) were prepared from the anterior part of the piriform cortex in an ice-cold artificial cerebro-spinal fluid (ACSF) solution saturated with 95% oxygen and 5% CO_2_. The ACSF solution contained (in mM) 125 NaCl, 25 NaNCO_3_, 25 Glucose, 3 KCl, 1.25 NaH2PO_4_, 2 CaCl_2_, 1 MgCl_2_ PH 7.4. The slices were incubated for 30 minutes at 37°c and kept at room temperature afterwards. During experiments, cells were visualized with a confocal scanning microscope equipped with infrared illumination and Dot gradient contrast video microscopy. Whole cell patch clamp recordings were performed using an Axon amplifier (Multi clamp). For patching, glass electrodes (6-8 MΩ) were made from thick-walled (0.25mm) borosilicate glass capillaries on a Flaming/Brown micropipette puller (P-97; Sutter Instrument). Intracellular pippet solution contained (in mM) 135 K+-gluconate, 4 KCl, 4 Mg-ATP, 10 Na2-phosphocreatine, 0.3 Na-GTP, 10 HEPES, 0.2 OGB-6F, 0.2 CF 633, and biocytin (0.2%;pH7.2).

Fluorescence confocal microscopy (Olympus FV1000) was performed on an upright BX61WI Olympus microscope equipped with a 60X (Olympus 0.9 NA) water objective. Neurons were filled with the calcium-sensitive dye OGB-6F (200 µM; Invitrogen) and CF 633 (200 µM; Biotium) to visualize the apical dendritic tree. Calcium transients were recorded in line-scan mode at 500 Hz.

All experiments were performed at 36° C.

All animal procedures were in accordance with guidelines established by the NIH on the care and use of animals in research and were confirmed by the Technion Institutional Animal Care and Use Committee.

### Focal stimulation

Focal synaptic stimulation, at apical dendrites of PCx pyramidal neurons was performed via a theta-glass (borosilicate; Hilgenberg) pipettes located in close proximity to the selected dendritic segment guided by the fluorescent image of the dendrite and the DIC image of the slice. The theta-stimulating electrodes were filled with CF-633 (Biotium; 0.2 mM). Current was delivered through the electrode (short burst of 3 pulses at 50 Hz), via stimulus isolator (ISO-Flex; AMPI). The efficacy and location of the stimulation was verified by simultaneous calcium imaging evoked by small EPSPs and their localization to a small segment of the stimulated dendrite.

### Glutamate uncaging

MNI-glutamate (Tocris, Bristol, UK) was delivered locally near by a dendritic region of interest using pressure ejection (5-10 mbar) from an electrode (2 µm in diameter) containing 5-10 mM caged glutamate. Electrodes were positioned 20-30 µm from the dendrite of interest and caged glutamate was photolyzed by a 1 ms laser pulse (375 nm Excelsior, Spectra Physics) using the point scan mode (Olympus FV1000). Simultanouse calcium imaging was performed from the uncaged dendritic region.

### Drug application

In part of the experiments, gamma-aminobutyric acid (GABAA) (1 µM bicuculline; Sigma) was added to the ACSF perfusion solution. In some experiments as indicated in the text, a cocktail of calcium channels blockers was added to the ACSF solution containing w-agatoxin 0.5 µM (P/Q type calcium channel blocker), conotoxin-GVIA 5µM (N type calcium channel blocker), SNX 482 200nM (R type calcium channel blocker) and nifedipine 10µM (L type calcium channel blocker). Sodium channel blocker TTX 1 µm was applied to the ACSF solution. NMDA-R antagonist APV (50 µM, Tocris Bioscience) was added to the ACSF solution. In some experiments the NMDAR channel blocker MK801 (1 mM) was addedd to the intracellular solution, and the tip of the electrode was backfilled with control intracellular pipette solution.

### Statistcal procedure

The sample size was chosen based on standards used in the field as well as our own vast experience with similar experimental paradigms. Importantly most of our experiments involve examining a variable on the same neuron and thus the sources of variability are smaller in these type of experiments.

Analysis was done with IgorPro (5.01; WaveMetrics), Exel and Clampfit (Molecular Devices) commercial softwares. Data are presented as mean ± SEM. For testing statistical significance we used two-tailed paired Student’s t test. No statistical methods were used to predetermine sample sizes. Our sample sizes are similar to those reported in previous publications. Neurons were excluded in case the viability of the cell was compromised as monitored by resting membrane potential and shape and amplitude of action potentials evoked by current injection. Average resting membrane potential was −80.1 ± 1.43 mV. In addition, we excluded neurons in which the quality of the recordings deteriorated as measured by the access resistance.

### Modeling

The simulations were conducted in a compartmental model using the NEURON 7.4 simulation platform. Three pyramidal cells were reconstructed from z-stacks of fluorescently labeled neurons using Simple Neurite Tracer (ImageJ, ^61^). The cells were subdivided into 493-544 compartments, with a maximum length of 19 µm. The soma area was 829 µm^2^, the total dendritic length was 2238-2516 µm^2^ ^44^. The resting membrane potential was −70 mV; the membrane resistance was 25,000 Ω·cm^2^; the axial resistance was 100 Ω·cm and the membrane capacitance was set to 1 µF/µm^2^. The simulations that included sodium and potassium voltage-gated currents used the Hodgkin-Huxley kinetics formalism. Specifically, fast sodium channels (reversal potential = 50 mV, gNa = 1000 mS/cm^2^), and delayed rectifier and slow non-inactivating potassium channels (reversal potential = −87 mV, gKdr = 500 mS/cm^2^, gKs = 20 mS/cm^2^) were used to allow for spike generation and adaptation, respectively ^27^.

To study the synaptic integration in PCx pyramidal neurons, we modeled physiologically realistic patterns of synaptic bombardment. We distributed the excitatory synapses from LOT presynaptic cells in the PCx pyramidal neurons according to known anatomical and physiological properties ^4,9,35,44^. The firing rates and the number of spikes for individual LOT presynaptic cells were drawn from random distributions that matched the known in-vivo firing properties of Mitral / PCx pyramidal cells ^9,18,41,42^. The stimulation intensity of a single presynaptic cell was set to produce an EPSC of 30 ± 19 pA, corresponding to a 1.4 ± 0.82 mV somatic EPSP (Fig S3 B; ^62^). Inhibitory inputs were randomly placed either in the LOT recipient band, even with the LOT excitation, or in the proximal apical dendritic region within 100 µm of the soma ^47^. For a clustered synaptic distribution, all excitatory inputs were placed on a single distal dendrite. In the dispersed distribution, excitatory inputs were allowed to target any LOT-recipient branch. To model intermediate clustering levels, we divided the excitatory input into two pools, one clustered and the second dispersed, and changed the proportion between the number of inputs in each pool. Presynaptic neurons were represented by NetStim processes that generated temporal triggers for synaptic activation. Each presynaptic cell gave rise to a single synapse on the modeled cell. Synaptic inputs were driven by a unique spike train for each presynaptic cell, which was generated by setting the ‘noise’ parameter of the NetStim process to 0.5. Excitatory spike trains began at simulation time of 100 ± 6 ms, and inhibitory inputs followed 10 ms later (110 ± 6ms) ^18,44^. The ISI and the number of presynaptic action potentials in the excitatory / inhibitory presynaptic populations were described by normal distributions (mean ± SD) of 8 ± 5 ms and 5 ± 3 / 10 ± 3 respectively ^9,18,45^.

Excitatory postsynaptic synapses contained AMPA-Rs and NMDA-Rs. Inhibition was mediated by GABA-A synaptic currents. GABA-A currents had an instantaneous rise time, a decay time of 7 ms, unitary conductance of 2 nS and reversal potential of −70 mV. All excitatory inputs reversed at 0 mV. AMPA-R currents had an instantaneous rise time and a decay time of 1.5 ms. The average unitary AMPA-R conductance was 1 nS ^26^. NMDA-R currents had a rise time of 2 ms and a decay time of 80 ms, and the average NMDA-R conductance was 2 nS. The NMDA-R conductance voltage dependence was modeled as follows: gNMDA= 1/(1+0.25·exp(−0.08·V_m_)) where V_m_ is the local membrane potential. In some simulations we canceled out the voltage dependence of the NMDA-R current by setting the V_m_ to −70 mV for the whole duration of the simulation ^27^. Presynaptic vesicular release was explicitly modeled; each synapse was assumed to contain 5 vesicles, each with an independent release probability (Pr) of 0.1. The presynaptic pool was replenished with a rate of 100 sec^−1^ ^63^).

